# Role of Mcpip1 in obesity-induced hepatic steatosis as determined by myeloid and liver-specific conditional knockouts

**DOI:** 10.1101/2020.09.17.301531

**Authors:** Natalia Pydyn, Dariusz Żurawek, Joanna Kozieł, Edyta Kuś, Kamila Wojnar-Lason, Agnieszka Jasztal, Mingui Fu, Jolanta Jura, Jerzy Kotlinowski

## Abstract

Monocyte chemoattractant protein-induced protein 1 (MCPIP1, *alias* Regnase1) is a negative regulator of inflammation, acting through cleavage of transcripts coding for proinflammatory cytokines and by inhibition of NFκB activity. Moreover, it was demonstrated, that MCPIP1 regulates lipid metabolism both in adipose tissue and hepatocytes. In this study, we investigated the effects of tissue-specific Mcpip1 deletion on the regulation of hepatic metabolism and development of non-alcoholic fatty liver disease (NAFLD).

We used knock-in control Mcpip1^fl/fl^ mice and animals with deletion of Mcpip1 in myeloid leukocytes (Mcpip1^fl/fl^LysM^Cre^) and in hepatocytes (Mcpip1^fl/fl^Alb^Cre^), which were fed chow or a high-fat diet (HFD) for 12 weeks. Mcpip1^fl/fl^LysM^Cre^ mice were fed a chow diet were characterized by a significantly reduced hepatic expression of genes regulating lipid and glucose metabolism, which subsequently resulted in hypoglycemia and dyslipidemia. These animals also displayed systemic inflammation, demonstrated by increased concentrations of cytokines in the plasma. On the other hand, there were no significant changes in phenotype in Mcpip1^fl/fl^Alb^Cre^ mice. Although we detected a reduced hepatic expression of genes regulating glucose metabolism and β-oxidation in these mice, they remained asymptomatic. Upon feeding them a HFD, Mcpip1^fl/fl^LysM^Cre^ mice did not develop obesity, glucose intolerance, nor hepatic steatosis, but were characterized by hypoglycemia and dyslipidemia, along with proinflammatory phenotype with symptoms of cachexia. Mcpip1^fl/fl^Alb^Cre^ animals, following a HFD, became hypercholesterolemic, but accumulated lipids in the liver at the same level as Mcpip1^fl/fl^ mice, and no changes in the level of soluble factors tested in the plasma were detected.

In conclusion, we have demonstrated that Mcpip1 protein plays an important role in the liver homeostasis. Depletion of Mcpip1 in myeloid leukocytes, followed by systemic inflammation, has a more pronounced effect on controlling liver metabolism and homeostasis than the depletion of Mcpip1 in hepatocytes.

## Introduction

Monocyte chemoattractant protein-induced protein 1 (MCPIP1, alias Regnase1), encoded by the *ZC3H12A* gene, is a RNase that degrades mRNAs, pre-miRNAs, and viral RNAs [1]. MCPIP1 functions as a negative regulator of inflammation, by direct cleaving of transcripts coding for proinflammatory cytokines like IL-1β, IL-6, IL-8 and IL-12 [2, 3, 4, 5], and by indirect inhibition of NFκB activity [6]. Apart from immunomodulatory effects, MCPIP1 activity was shown to regulate many biological processes, such as cell differentiation, angiogenesis, proliferation, and adipogenesis [7, 8, 9]. Adipogenesis of 3T3-L1 cells is inhibited *via* the degradation of C/EBPβ and selected pre-miRNAs by MCPIP1 [9, 10]. Additionally, the MCPIP1 level in human adipose tissue inversely correlates with patients’ BMI [11]. Similarly to humans, the protein level of Mcpip1 is reduced in hepatocytes and in subcutaneous adipose tissue of mice with NAFLD, in comparison to control animals [12]. Despite the results mentioned above, the functions of Mcpip1 in liver metabolism and NAFLD progression remain obscure.

Previous studies have shown that Mcpip1-deficient mice spontaneously develop systemic inflammatory response, leading to splenomegaly, lymphadenopathy, hyperimmunoglobulinemia, and ultimately death within 12 weeks [3, 13]. Similarly, a hematopoietic deficiency of Mcpip1 resulted in severe systemic and multi-organ inflammation [14]. It was also shown that overexpression of Mcpip1 reduces liver injury in septic mice, by inhibiting inflammatory reaction in macrophages [15]. Additionally, Mcpip1 ameliorates liver damage, reduces inflammation, and promotes tissue regeneration in the hepatic ischemia/reperfusion injury model [16].

NAFLD describes a wide range of liver conditions affecting people who drink little to no alcohol. It is a chronic and progressive disease, characterized in the first stages by an excessive accumulation of triglycerides in hepatocytes [17]. Untreated NAFLD can progress from simple steatosis, to more severe non-alcoholic steatohepatitis (NASH), which is additionally characterized by liver inflammation and fibrosis [18]. Ultimately, NASH can further progress to cirrhosis and/or hepatocellular carcinoma, which over time might require liver transplantation [19].

In present study, we aimed to examine the contribution of Mcpip1 to liver metabolism and the pathogenesis of NAFLD. Specifically, we used Cre-LoxP technology to delete *Zc3h12a* gene encoding Mcpip1 in myeloid leukocytes (Mcpip1^fl/fl^LysM^Cre^), and in hepatocytes (Mcpip1^fl/fl^Alb^Cre^). Myeloid cells are represented in the liver mostly by Kupffer cells, that make up 80% of whole-body macrophages, which is the largest population of tissue macrophages in solid organs [20]. Crosstalk between Kupffer cells and hepatocytes, mediated for example by IL-1β, was shown to promote hepatic triglyceride storage through IL-1β-dependent suppression of PPARα activity, and subsequent inhibition of β-oxidation [21]. Also, blood monocytes infiltrating the liver during NAFLD are involved in its progression through the production of a vast array of proinflammatory mediators [22]. In our study, by comparing Mcpip1^fl/fl^LysM^Cre^ and Mcpip1^fl/fl^Alb^Cre^ animal models, we were able to analyze the tissue-specific effects of Mcpip1 on liver metabolism and NAFLD development. Our results provide comprehensive data on the role of the Mcpip1 expression in the myeloid leukocytes and hepatocytes in the regulation of metabolism and sterile inflammation.

## Materials and Methods

### Animals, diets, and genotyping

To obtain myeloid or hepatocyte-specific knockout of Mcpip1 protein, we used Mcpip1^fl/fl^ mice with two loxp sites flanking the two sides of exon 3 of the *Zc3h12a* gene encoding Mcpip1 [23]. Next, these mice were crossed with myeloid cells (LysM^CreTg/Wt^, Jackson Laboratory) or hepatocytes expressing Cre transgenic mice (Alb^CreTg/Wt^, Jackson Laboratory). In the second round of crosses, Mcpip1^fl/fl^ females were crossed with Mcpip1^fl/fl^ LysM^CreTg/Wt^ males to obtain both control Mcpip1^fl/fl^ LysM^CreWt/Wt^ animals (designed as Mcpip1^fl/fl^) and Mcpip1^fl/fl^ LysM^CreTg/Wt^ knockouts (designed as Mcpip1^fl/fl^LysM^Cre^). Similarly, Mcpip1^fl/fl^ females were crossed with Mcpip1^fl/fl^Alb^CreTg/Wt^ males to obtain controls (Mcpip1^fl/fl^) and knockouts (Mcpip1^fl/fl^Alb^Cre^). All mice had C57BL/6N background, and were fed *ad libitum* for 12 weeks with a control diet or HFD, 60% kcal from fat) (ZooLab), starting from the age of 10 weeks. Animals were housed under SPF conditions in ventilated cages in a temperature-controlled environment with a 14/10◻h light/dark cycle. To measure the amount of consumed food, the mice were housed in metabolic cages. For genotyping, DNA was extracted from tail tissue with a KAPA Mouse Genotyping Kit (KAPA Biosystems) according to the manufacturer’s instructions. Sequences of primers (Sigma) used for genotyping are listed in Table S1 in Supplementary Material. All animal procedures were conducted in accordance with the Guide for the Care and Use of Laboratory Animals (Directive 2010/63/EU of the European Parliament) and carried out under a license from the 2^nd^ Local Institutional Animal Care and Use Committee in Kraków (study no. 106/2017).

### Blood and tissue samples

The mice were fasted for 4 h, after which their blood was collected from the orbital sinus after isoflurane anesthesia into EDTA containing tubes, and then the animals were euthanized. Plasma was prepared by centrifugation (1000g, 10 min) and stored at −80°C. Following euthanasia by cervical dislocation, organs were isolated and prepared for histological analysis, or snap-frozen in liquid nitrogen and stored at −80°C.

### Plasma analysis

Murine plasma was used for the subsequent analysis: total cholesterol, HDL-cholesterol (HDL), LDL-cholesterol (LDL), glucose, triglycerides, alanine aminotransferase (ALT), aspartate aminotransferase (AST), and lactate dehydrogenase (LDH). These parameters were measured using an automatic biochemical analyzer, Pentra 400 (Horiba), by an enzymatic photometric method according to the vendor’s instructions.

### Glucose tolerance test

A glucose tolerance test (GTT) was performed on mice aged 10 and 22 weeks. Mice were fasted for 4 hours, after which they were intraperitoneally injected with saturated glucose solution (2◻g/kg body weight). Blood was collected from the tail vein before (0◻min) and at 15, 30, 45, 60, 120◻min after glucose application. Glucose concentrations were measured by an Accu-Check glucometer (Roche Diagnostics). The area under the curve (AUC) of blood glucose concentration was calculated geometrically by applying a trapezoidal method [24].

### Histological analysis

Livers were fixed in 4% paraformaldehyde formalin solution and divided into two parts. One piece was submerged in a 30% sucrose solution overnight for cryoprotection, and then frozen in Tissue-Tek® OCT medium at −80◻°C. Frozen sections were stained using an Oil Red-O (ORO) method to analyze fat accumulation, and imaged under 100◻x magnification. 10 images of each section were randomly collected and analyzed by the Columbus v.2.4.2 software (Perkin Elmer) with an algorithm adapted for Oil Red-O stained sections. The second piece was processed using standard paraffin procedures, and 5-μm paraffin tissue sections were stained with Hematoxylin and Eosin (H&E) for general histology and Picro Sirius Red (PSR) for collagen deposition, and then visualized using a standard light microscope (Olympus BX51; Olympus Corporation).

### Luminex assay

Magnetic Luminex Assay (R&D Systems) was performed on plasma samples, according to the manufacturer’s instructions. All samples were analyzed in duplicate on a MAGPIXs (Luminex, Texas, USA). Median fluorescence intensity (MFI) data, obtained using a weighted 5-parameter logistic curve-fitting method, was used to calculate analyte concentrations.

### RNA isolation and RT-PCR

Total RNA was isolated from liver tissue using a modified guanidinium isothiocyanate method [25]. Briefly, fragments of frozen liver tissue were lysed in fenozol (A&A Biotechnology), and after addition of chloroform, the aqueous phase containing RNA was collected into new eppendorf tubes. Next, RNA was precipitated, centrifuged, washed in ethanol and finally dissolved in nuclease-free water. The concentration of RNA was assessed by a NanoDrop 1000 Spectrophotometer (Thermo Fisher Scientific). Reverse transcription was performed using 1 μg of total RNA, oligo(dT) 15primer (Promega) and M-MLV reverse transcriptase (Promega). Real-time PCR was carried out using Sybr Green Master Mix (A&A Biotechnology) and QuantStudio Real-Time PCR System (Applied Biosystems). Gene expression was normalized to elongation factor-2 (EF2), after which the relative level of transcripts was quantified by a 2^−ΔΔCT^ method. Sequences of primers (Genomed/Sigma) and annealing temperatures are listed in Table S2 in Supplementary Material. The primers were designed to cross an exon-exon boundary to allow the specific amplification of cDNA targets.

### Measurement of body temperature

Mice body temperature was measured by rectal probing (physiological monitoring unit THM150, Visualsonics). The probe was inserted for 2 cm into the rectum and held in position for 10 seconds before temperature was determined. Measurements were conducted for five days at exactly the same time of day.

### Open field test

All mice were handled for three consecutive days prior to the start of behavioral measurements. On the day of the behavioral experiment, the mice were brought in their home cages to the testing room and left for one hour to acclimate. General locomotor activity and movement velocity were monitored in an open field in arena cages (40 x 40 x 38 cm) made from transparent plexiglass, equipped with an array of infrared photo beams and software for automatic tracking of mouse ambulation (Activity Monitor 7, Med Associates inc. St. Albans, VT, USA). The open field arena was illuminated with white light at a level of 150 lux. The duration of each behavioral session was 30 min for each animal. After each trial, the cages were wiped with 70% ethanol and left to dry. To eliminate the effect of circadian rhythm on locomotor activity, all mice were subjected to the open field test between 6 and 9 pm.

### Statistical analysis

Results are expressed as mean ± SEM. The exact number of experiments or animals used is indicated in the figure legends. One-way ANOVA with Tukey’s post hoc test was applied for comparison of multiple groups. To analyze the effects of two variable factors, a two-way ANOVA was used. The p values are marked with an asterisks in the charts (* p<0.05; ** p<0.01; *** p<0.001).

## Results

### Deletion of Mcpip1 in myeloid cells, but not in hepatocytes, results in dyslipidemia

Control Mcpip1^fl/fl^ animals used in this study were born in both Mcpip1^fl/fl^LysM^Cre^ and Mcpip1^f/fl^Alb^Cre+^ mice colonies. In the first analysis, we tested whether there is a difference between these animals. As compared in figure S1, there were no changes in mass, blood biochemistry, gene expression, nor in plasma cytokines concentrations in both Mcpip1^fl/fl^ control cohorts (Fig. S1). Thus, in the next figures, all data are presented with one control group.

Starting from the 7^th^ week of the experiment, Mcpip1^fl/fl^LysM^Cre^ mice were characterized by reduced body weight, although they consumed an equal amount of food to Mcpip1^fl/fl^ (Fig. 1A,B, S2A). At the age of 22 weeks, Mcpip1^fl/fl^LysM^Cre^ exhibited hepatomegaly and significantly lower cholesterol, HDL levels with concomitant increased concentration of LDL, when compared to control Mcpip1^fl/fl^ mice (Fig. S2B,C, 1C-E). We did not observe differences in the concentration of triglycerides, nor activity of ALT, AST and LDH in Mcpip1^fl/fl^LysM^Cre^ animals (Fig. 1F,G S2D,E). Hypoglycemia of Mcpip1^fl/fl^LysM^Cre^ mice was demonstrated by glucose tolerance test (Fig. 1H,I). Additionally, all measured analytes did not differ between control and Mcpip1^fl/fl^Alb^Cre^ mice, however Mcpip1 depletion in hepatocytes resulted in hepatomegaly (Fig. 1C-I, S2B-E).

**Figure 1:**
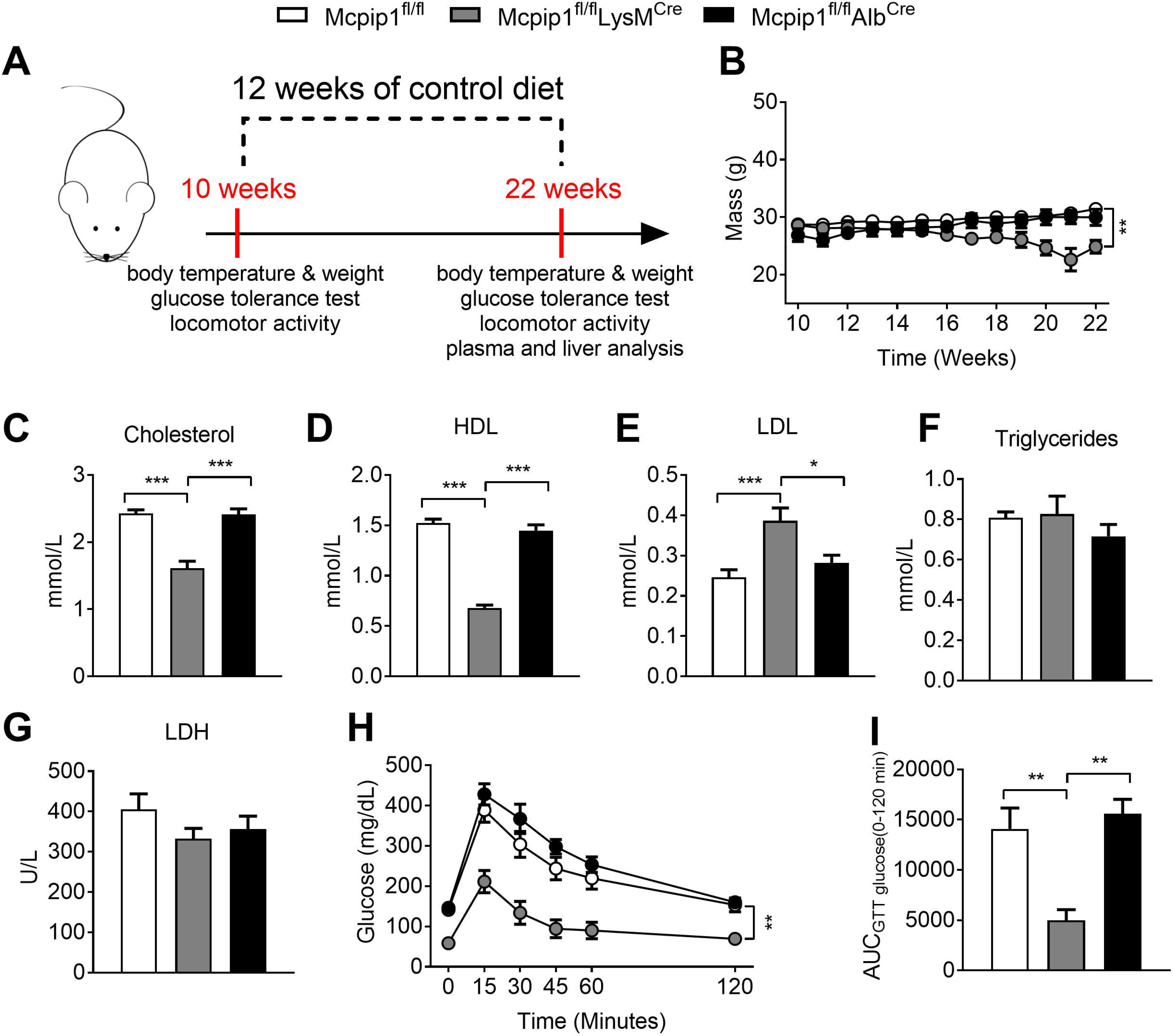
Mcpip1^fl/fl^LysM^Cre^ mice are characterized by reduced body weight, dyslipidemia and hypoglycemia. Mcpip1^fl/fl^, Mcpip1^fl/fl^LysM^Cre^ and Mcpip1^fl/fl^Alb^Cre^ mice were analyzed at the age of 10 and 22 weeks (A). (B) Body weight measurements of Mcpip1^fl/fl^, Mcpip1^fl/fl^LysM^Cre^ and Mcpip1^fl/fl^Alb^Cre^ mice fed chow diet. Plasma analysis of 22-week old mice: (C) total cholesterol; (D) HDL; (E) LDL; (F) Triglycerides; (G) LDH levels. Glucose metabolism was tested after intraperitoneal injection of 2 g/kg of glucose by (H) glucose tolerance test and by (I) calculations of the area under the curve (AUC). For Mcpip1^fl/fl^ n=10 (Fig. 1B-G) and n=8 (Fig. 1H,I), for Mcpip1^fl/fl^LysM^Cre^ n=6 (Fig. 1B-G) and n=5 (Fig. 1H,I), for Mcpip1^fl/fl^Alb^Cre^ n=8 (Fig. 1B-G) and n=6 (Fig. 1H,I). The graphs show means ± SEM; * p<0.05; ** p<0.01; *** p<0.001.

### Mcpip1 depletion leads to liver pathology

Histological analysis of livers collected from Mcpip1^fl/fl^LysM^Cre^ mice showed pathology of the bile ducts, while livers of Mcpip1^fl/fl^ mice did not show any pathological changes (Fig. 2). The observed changes include hyperplasia and hypertrophy of cholangiocytes, as well as changes of a dysplastic nature. The most intense changes were observed in the large interstitial bile ducts were the pathology of cholangiocytes also included hyalinization and ultimately cytoplasmic degranulation and cell disintegration. Moreover, observed changes were accompanied by intense inflammatory infiltration and local fibrosis without penetration into the liver parenchyma (Fig. 2A). Necrosis of single hepatocytes were found only in areas of inflammation associated with extreme degenerative changes in the bile ducts. Additionally, Mcpip1^fl/fl^LysM^Cre^ mice were characterized by an increased extramedullary hematopoiesis in the liver. Hematopoietic foci were located mainly in the walls of the sub-lobular veins, significantly thickening them and narrowing the lumen of these vessels. The local fibrosis limited to the area of hematopoietic islets were also observed (Fig. 2B). Histological analysis of livers collected from Mcpip1^fl/fl^Alb^Cre^ mice revealed very intense pathology of the intrahepatic bile ducts. These mice displayed hyperplasia of small bile ducts with accompanying hypertrophy of cholangiocytes, disturbance of their tubular structure and lumen obstruction (Fig. 2A). Comparison of livers collected form Mcpip1^fl/fl^Alb^Cre^ mice with control ones showed also infiltration of mononuclear cells and enhanced deposition of collagen. Significant fibrosis was detected within the portal space, around bile ducts, and to lower extend around the central veins and in the remaining area of the parenchyma (Fig. 2B).

**Figure 2:**
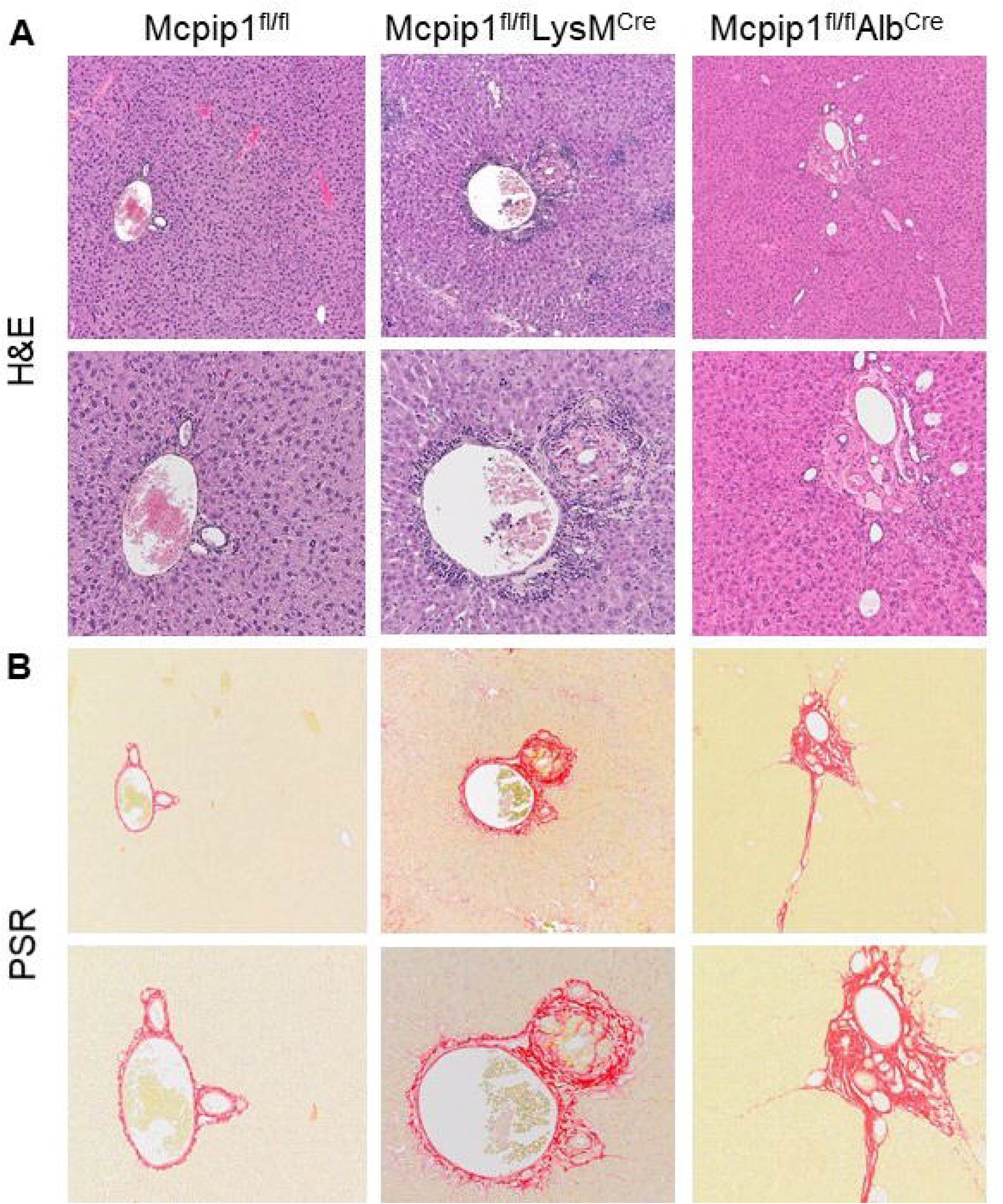
Mcpip1 depletion leads to liver pathology. Representative liver (A) hematoxylin & eosin (H&E), and (B) Picro Sirius Red (PSR) stainings. 100x and 200x magnification.

### Myeloid and hepatic Mcpip1 deletion alters expression of genes regulating glucose metabolism

In order to identify the molecular mechanism responsible for the development of hypoglycemia and dyslipidemia in Mcpip1^fl/fl^LysM^Cre^ animals, we analyzed a hepatic gene expression. The analysis revealed that Mcpip1^LysKO^ exhibit a significantly reduced expression of *G6pc* (regulating gluconeogenesis), *Irs1, Irs2* (response to insulin) and *Slc2a2* (glucose bidirectional transport) in livers (Fig. 3A). Changes in the livers of Mcpip1^fl/fl^Alb^Cre^ mice were not as significant as in the case of Mcpip1^LysKO^, we detected lower expression of *Pck1, G6pc* and *Irs1* in comparison to Mcpip1^fl/fl^ mice (Fig. 3A).

**Figure 3:**
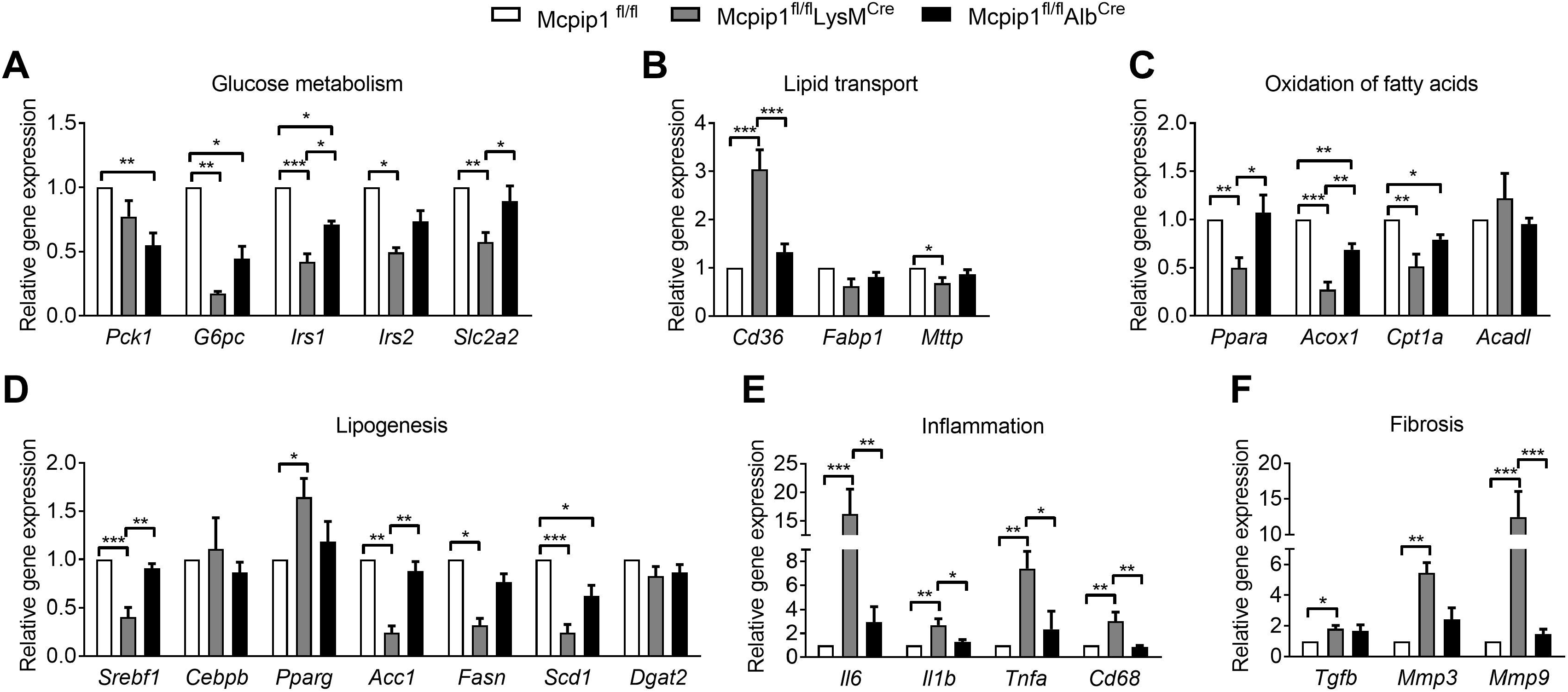
Expression of Mcpip1 is essential for proper liver metabolism. Expression of genes coding for proteins involved in (A) glucose metabolism; (B) lipid transport; (C) oxidation of fatty acids; (D) lipogenesis; (E) inflammation and (F) fibrosis. For Mcpip1^fl/fl^ n=8, for Mcpip1^fl/fl^LysM^Cre^ n=6, for Mcpip1^fl/fl^Alb^Cre^ n=9. The graphs show means ± SEM; * p<0.05; ** p<0.01; *** p<0.001.

### Mcpip1 depletion disturbs lipid metabolism in the livers of Mcpip1^fl/fl^LysM^Cre^ and Mcpip1^fl/fl^Alb^Cre^ mice

Besides regulating glucose metabolism, the liver is also a prominent regulator of lipid metabolism, due to the ability to perform lipoprotein synthesis and lipid storage [26]. Therefore, in the following set of experiments, we have evaluated the expression of key genes related to lipid transport, lipogenesis, and fatty acids oxidation. As shown in Figure 3B-D, more significant changes in hepatic lipid metabolism were detected in Mcpip1^fl/fl^LysM^Cre^ than in Mcpip1^fl/fl^Alb^Cre^ mice (Fig. 3B-D). The expression of *Cd36* was increased, while *Mttp* was downregulated in the livers of Mcpip1^fl/fl^LysM^Cre^ mice (Fig. 3B). Furthermore, the expression of *Ppara* was reduced in the hepatic tissue of Mcpip1^fl/fl^LysM^Cre^ mice, together with its downstream targets *Acox1* and *Cpt1a,* but not *Acadl* (Fig. 3C). Among the three master regulators of lipogenesis (*Srebf1, Cebpb, Pparg*) [27,28], we detected a lower expression of *Srebf1* in samples from Mcpip1^fl/fl^LysM^Cre^ mice, together with the downregulation of SREBP1c-regulated genes, *Acc1, Fasn* and *Scd1* (Fig. 3D). In livers isolated from Mcpip1^fl/fl^Alb^Cre^ animals, there were no differences in the expression of genes regulating lipid transport (*Cd36*, *Fabp1*, *Mttp*), but the amount of *Acox1*, *Cpt1a* and *Scd1* transcripts was significantly reduced in comparison to control Mcpip1^fl/fl^ mice (Fig. 3B-D).

### Deletion of Mcpip1 in myeloid cells, but not in hepatocytes, induces systemic inflammation

To further characterize the phenotype of both Mcpip1 knockout strains, we performed a Luminex Assay. Out of 45 analytes, 14 were not detected in all samples, whereas an expression of 20 was significantly changed between Mcpip1^fl/fl^LysM^Cre^ and control mice (Table 1). The level of cytokines and its receptors (Tnf-alpha, Il-6 R alpha, Tnf RI, Tnf RII) was significantly upregulated in Mcpip1^fl/fl^LysM^Cre^ in comparison to Mcpip1^fl/fl^ mice. Cytokines that are chemotactic among other monocytes (Ccl8/Mcp2, Ccl7/Marc, Ccl3/Mip-1 alpha) dendritic cells, T and B cells (Ccl19/Mip-3 beta) lymphocytes, and eosinophils (Ccl12/Mcp-5), were significantly induced in Mcpip1^fl/fl^LysM^Cre^ animals. Concentrations of other inflammation markers, such as Syndecan-1/CD138, Chi3-L1, and p-Selectin, were also increased in Mcpip1^fl/fl^LysM^Cre^ animals compared to control mice (Table 1). In myeloid Mcpip1 knockouts, we detected a reduced amount of three proteins, namely Vegf-R2, Thrombospondin 4, and endoglin. Surprisingly, concentrations of all tested analytes were not changed between Mcpip1^fl/fl^ and Mcpip1^fl/fl^Alb^Cre^ mice, indicating a key immunomodulatory role of myeloid Mcpip1.

**Table 1:**
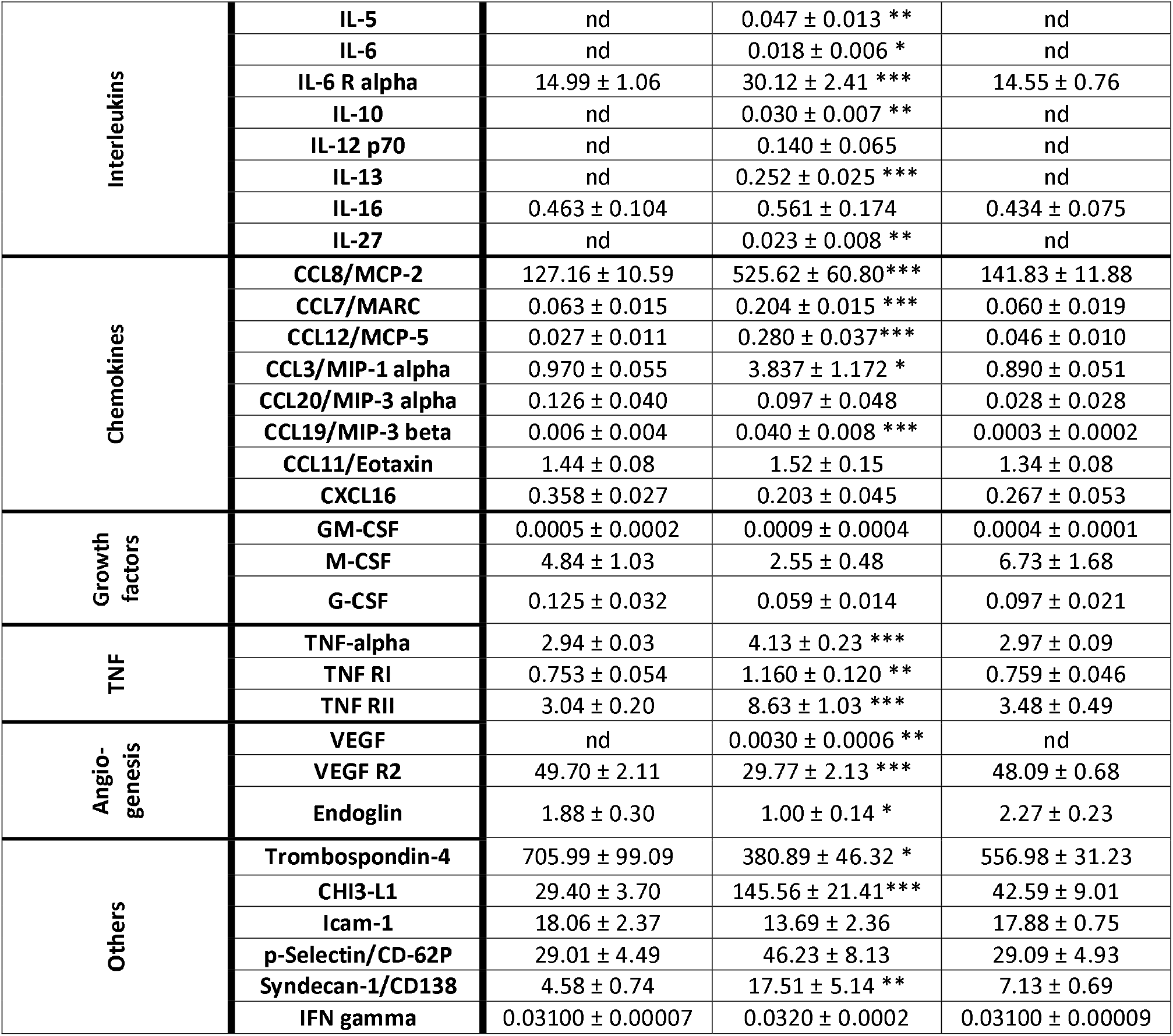
Immune profile of plasma samples from 22 week-old mice determined by Luminex Assay. Out of 45 analytes, 14 has not been detected, namely: IL-2, IL-1β, IL-3, IL-7, CCL4/MIP-1 beta, CCL2/MCP1, RAGE, IL17e/IL25, IL17/IL17a, IL1a, IL-4, IL-33, CCL5/RANTES, FGF-basic. For Mcpip1^fl/fl^ n=6, for Mcpip1^fl/fl^LysM^Cre^ n=7, for Mcpip1^fl/fl^Alb^Cre^ n=7. The table shows means ± SEM; * p<0.05; ** p<0.01; *** p<0.001, nd – not detected.

Systemic inflammation in Mcpip1^LysKO^ mice was followed by a proinflammatory hepatic gene expression pattern. Expression of *Il6, Il1b*, as well as *Tnfa* and *Cd68* was significantly upregulated in the livers of Mcpip1^fl/fl^LysM^Cre^ in comparison to Mcpip1^fl/fl^ mice. However, there were no changes between Mcpip1^fl/fl^ and Mcpip1^fl/fl^Alb^Cre^ samples (Fig. 3E). Moreover, we have found a significantly increased expression of profibrotic genes – *Tgfb*, *Mmp3* and *Mmp9* – in the livers of Mcpip1^fl/fl^LysM^Cre^. On the other hand, Mcpip1^fl/fl^Alb^Cre^ mice were characterized by slightly, but not significantly higher expression of these selected genes (Fig. 3F).

### HFD does not induce obesity and hepatic steatosis in Mcpip1^fl/fl^LysM^Cre^ mice

In agreement with literature data, HFD feeding increased body mass and liver steatosis of Mcpip1^fl/fl^ mice (Fig. 4A-C). However, starting from the third week of experiment, Mcpip1^fl/fl^LysM^Cre^ mice were characterized by significantly decreased body weight, that cannot be fully attributed to lower food consumption (Fig. 4A, S2F). Additionally, Mcpip1^fl/fl^LysM^Cre^ mice, after 12 weeks of HFD, were characterized by very limited liver steatosis (3% vs 37% in Mcpip1^fl/fl^) and hepatomegaly (Fig. 4B,C, S2G,H). On the other hand, the depletion of Mcpip1 in Mcpip1^fl/fl^Alb^Cre^ mice did not cause any changes in body mass nor steatosis, but mice had hepatomegaly (Fig. 4A,B, S2G,H). However, liver of these animals had very pronounced collagen deposition accompanying hyperplasia of bile duct epithelial cell (Fig. 4C). Changes in lipid profile measured in plasma of both Mcpip1 knockouts, after 12 weeks of HFD, resembled results obtained in mice fed chow food: we detected low cholesterol and HDL concentration only in Mcpip1^fl/fl^LysM^Cre^ mice (Fig. 4D,E). 12 weeks of HFD increased the LDL level in Mcpip1^fl/fl^ in comparison to chow diet, blunting differences between these mice and myeloid knockouts (Fig. 4F). There were no changes in the LDL, triglycerides, ALT and AST in Mcpip1^fl/fl^LysM^Cre^ when compared to Mcpip1^fl/fl^ mice (Fig. 4F,G, S2I,J). Only the LDH level was raised in the plasma of Mcpip1^fl/fl^LysM^Cre^ mice (Fig. 4H). Mcpip1^fl/fl^Alb^Cre^ mice were characterized by increased cholesterol level, but all other analytes did not differ between these animals and Mcpip1^fl/fl^ (Fig. 4D-H, S2I,J). Finally, feeding with HFD led to hyperglycemia and an impaired glucose tolerance detected in all three strains in comparison to mice fed a chow diet (Fig. 4I,J). However, based on GTT test and fasting glucose measurement, Mcpip1^fl/fl^LysM^Cre^ animals were characterized by hypoglycemia, in comparison to Mcpip1^fl/fl^ counterparts (Fig. 4K,L).

**Figure 4:**
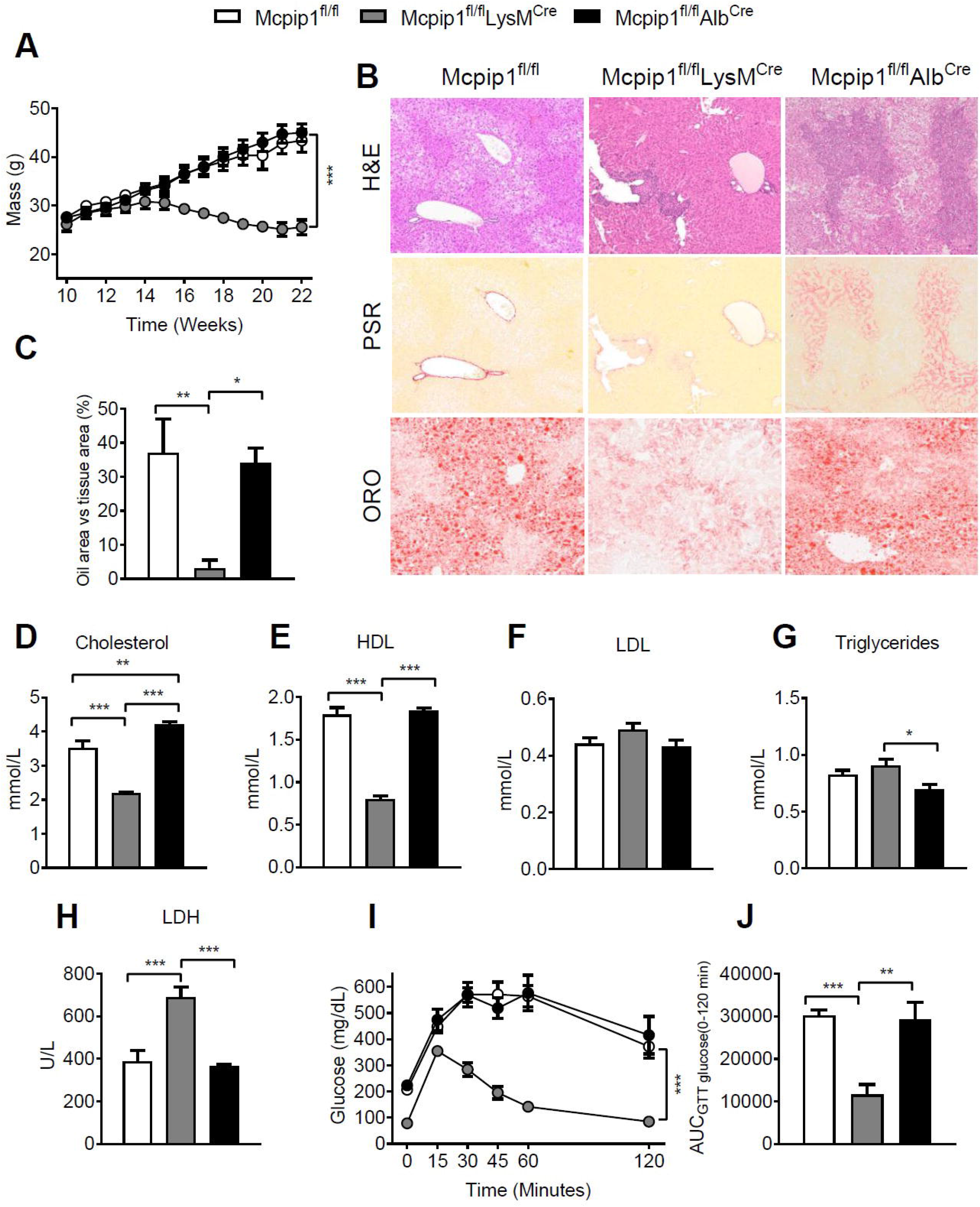
Mcpip1^fl/fl^LysM^Cre^ mice do not develop obesity and hepatic steatosis upon feeding with high fat diet. (A) Body weight measurements of Mcpip1^fl/fl^, Mcpip1^fl/fl^LysM^Cre^ and Mcpip1^fl/fl^Alb^Cre^ mice fed high fat diet for 12 weeks. (B) Representative liver hematoxylin & eosin (H&E), Oil Red O (ORO) and Picro Sirius Red (PSR) stainings; 100x magnification. (C) Quantitative analysis of Oil Red O-stained liver tissues; Plasma analysis of 22-week old mice: (D) Total cholesterol; (E) HDL; (F) LDL; (G) Triglycerides; (H) LDH levels. Glucose metabolism was tested after intraperitoneal injection of 2 g/kg of glucose by (I) glucose tolerance test and by (J) calculations of the AUC. For Mcpip1^fl/fl^ n=7, for Mcpip1^fl/fl^LysM^Cre^ n=6, for Mcpip1^fl/fl^Alb^Cre^ n=6. The graphs show means ± SEM; * p<0.05; ** p<0.01; *** p<0.001.

The profile of cytokines, chemokines, and growth factors in the plasma of HFD-fed mice showed similar tendencies to animals fed chow food (Table S2). Concentrations of proinflammatory mediators (interleukins, chemokines, soluble receptors, and ECM proteins) were significantly increased in Mcpip1^fl/fl^LysM^Cre^ mice compared to their Mcpip1^fl/fl^ counterparts (Table S2). Again, there were no differences between Mcpip1^fl/fl^Alb^Cre^ and Mcpip1^fl/fl^ mice. Interestingly, HFD feeding of Mcpip1^fl/fl^LysM^Cre^ mice further elevated the amount of MIP-1alpha, syndecan-1, and CHI3-L1, in comparison to animals on chow food (Table 1, S2).

### Mcpip1 deficiency changes hepatic mRNA expression profile in HFD-fed mice

The expression of key gluconeogenesis regulators (*Pck1, G6pc*) was not changed in myeloid knockouts, but impaired glucose metabolism in these mice is at least partially explained by a low level of *Irs1* and *Slc2a2* (Fig. 5A). Lipid metabolism was also altered in Mcpip1^fl/fl^LysM^Cre^ animals fed for 12 weeks with HFD, as manifested by the reduced expression of *Fabp1* and *Mttp* regulating lipid transport, and low levels of *Ppara*, *Acox1* and *Cpt1a* mRNAs, that are involved in beta-oxidation (Fig. 5B-C). Additionally, the expression of genes regulating lipogenesis (*Srebf1, Acc1, Fasn* and *Scd1*) was significantly reduced in Mcpip1^fl/fl^LysM^Cre^ mice (Fig. 5D). Additionally, there was a significant increase of proinflammatory and profibrotic transcripts in Mcpip1^fl/fl^LysM^Cre^ mice in comparison to their Mcpip1^fl/fl^ counterparts (Fig. 5E,F).

**Figure 5:**
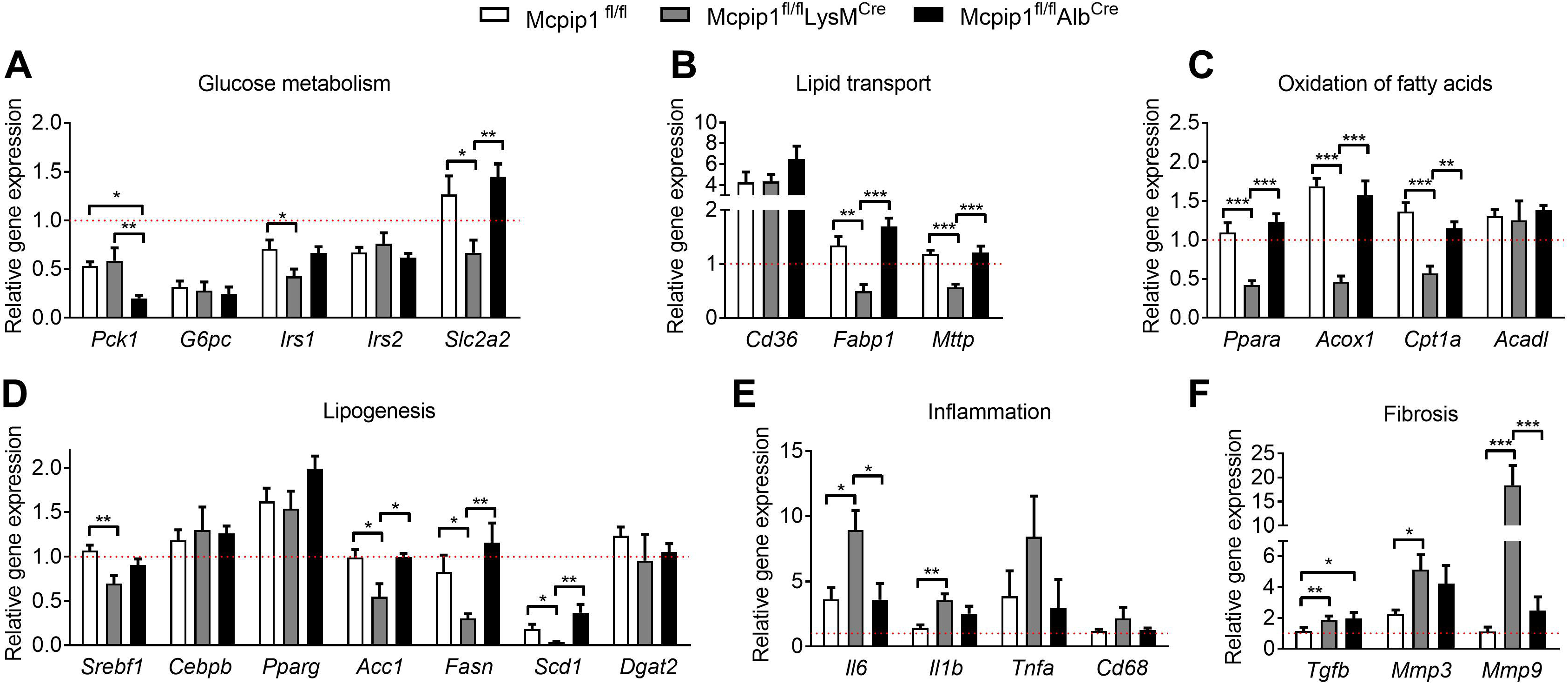
Disturbed hepatic gene expression profile is linked to lean phenotype of HFD-fed Mcpip1^fl/fl^LysM^Cre^ mice. Expression of genes coding for proteins involved in (A) glucose metabolism; (B) lipid transport (C) oxidation of fatty acids; (D) lipogenesis; (E) inflammation and (F) fibrosis. All results were calculated in comparison to appropriate gene in Mcpip1^fl/fl^ mice fed chow diet (ΔΔCt method). Dotted lines represent results obtained from Mcpip1^fl/fl^ mice fed chow diet. For Mcpip1^fl/fl^ n=7, for Mcpip1^fl/fl^LysM^Cre^ n=8, for Mcpip1^fl/fl^Alb^Cre^ n=9. The graphs show means ± SEM; * p<0.05; ** p<0.01; *** p<0.001.

HFD feeding for 12 weeks blunted differences between Mcpip1^fl/fl^Alb^Cre^ and Mcpip1^fl/fl^ mice. Out of 26 tested genes, only the level of *Pck1* was decreased, and *Tgfb* was increased in Mcpip1^fl/fl^Alb^Cre^ mice in comparison to controls (Fig. 5A,F). Overall changes in the hepatic gene expression pattern in HFD-fed mice – Mcpip1^fl/fl^LysM^Cre^ and Mcpip1^fl/fl^Alb^Cre^ knockouts vs. Mcpip1^fl/fl^ were very similar to mice fed chow food (Fig. 3A-F, 5A-F). Thus, our results indicate that the deletion of Mcpip1 protein in myeloid leukocytes has a more severe impact on metabolic regulatory function than hepatic Mcpip1 deficiency. Additionally, Mcpip1 acts independently from HFD-induced obesity effects on liver metabolic regulation.

### Myeloid Mcpip1 deletion impairs locomotor activity and causes hypothermia

Finally, it is well known that reduced locomotor activity is a major contributor to diet-induced obesity in mice [29]. That is why, in the last set of experiments, we investigated both average velocity and total traveled distance by 10 and 22-week-old Mcpip1^fl/fl^LysM^Cre^ and Mcpip1^fl/fl^Alb^Cre^ knockouts. Locomotor activity of all strains at 10 weeks was the same (Fig. 6A-B). However, 22-week-old Mcpip1^fl/fl^LysM^Cre^ mice were characterized by significantly reduced locomotor activity. Although the average velocity of Mcpip1^fl/fl^LysM^Cre^ was not changed, their total traveled distance was reduced by 70% in comparison to Mcpip1^fl/fl^ mice (Fig. 6A-B). These mice weighted on average 7 grams less than Mcpip1^fl/fl^ and Mcpip1^fl/fl^Alb^Cre^ mice, and were also characterized by hypothermia (35.5°C vs 37.6°C in controls) (Fig. 6C-D). In contrast, locomotor activity, weight and temperature in 22-week-old Mcpip1^fl/fl^Alb^Cre^ mice were not affected by Mcpip1 deletion (Fig. 6A-D).

**Figure 6:**
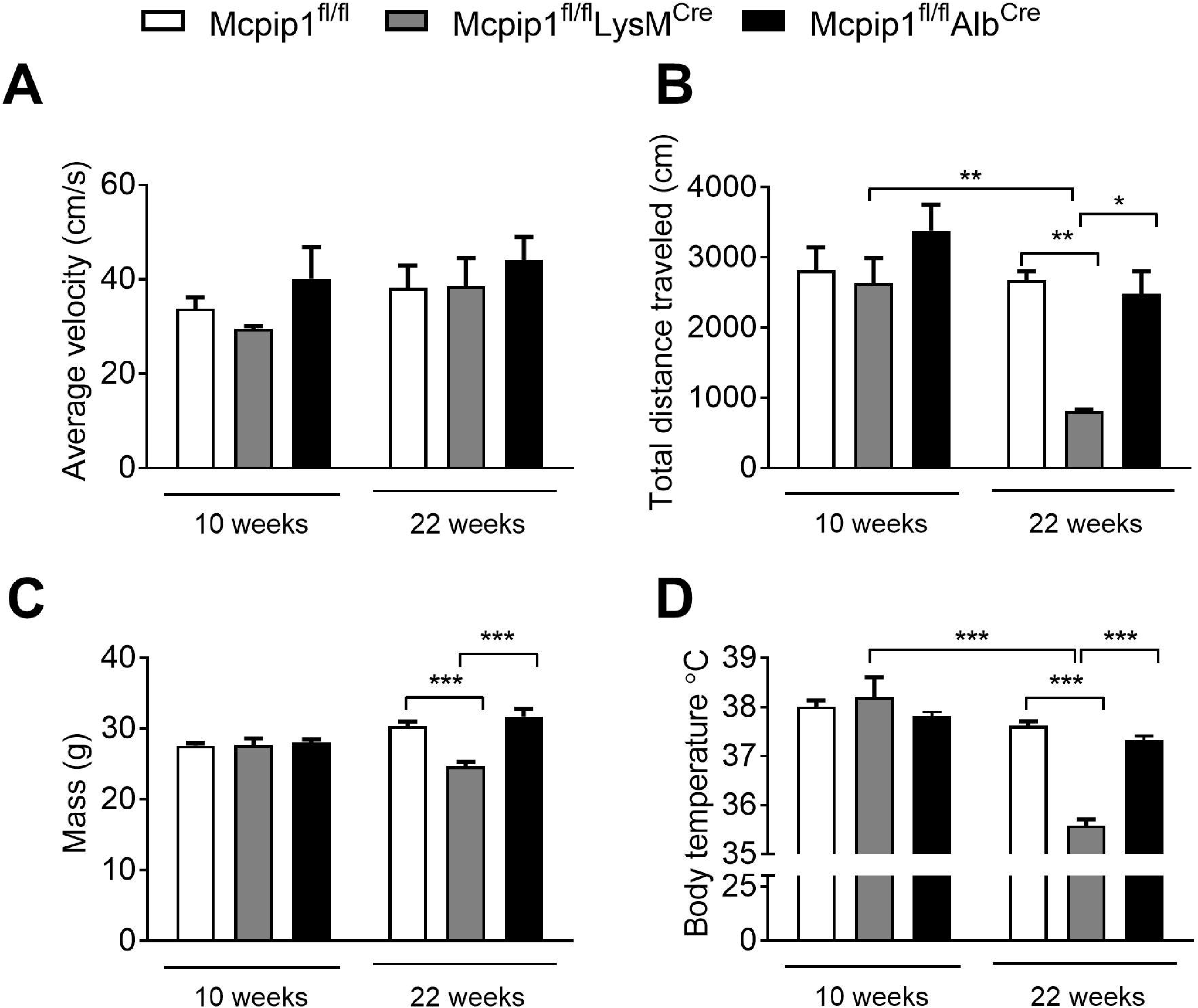
Myeloid Mcpip1 deletion impairs locomotor activity and causes hypothermia. Locomotor activity was measured at both time points by changes in (A) average velocity and (B) total distance traveled. (C) Body weight; and (D) body temperature. For Mcpip1^fl/fl^ n=10, for Mcpip1^fl/fl^LysM^Cre^ n=5, for Mcpip1^fl/fl^Alb^Cre^ n=6. The graphs show means ± SEM; * p<0.05; ** p<0.01; *** p<0.001.

## Discussion

We previously demonstrated that hepatic steatosis accompanying diet-induced obesity diminishes the amount of Mcpip1 protein in murine primary hepatocytes [12]. Similarly, the Mcpip1 protein level was reduced in murine and human adipose tissue collected from obese individuals [11, 12]. Additionally, Mcpip1 was shown to regulate lipid and glucose metabolism in hepatic and adipose tissue. Mechanistically, in hepatocytes Mcpip1 induces both the expression and activity of hepatic peroxisome proliferator-activated receptor gamma *via* the TXNIP/PGC-1α pathway [12]. On the other hand, in 3T3-L1, preadipocytes differentiated into adipocytes, and endogenous MCPIP1 is downregulated at early stages of differentiation, allowing fat accumulation by preadipocytes. Additionally, an overexpression of MCPIP1 in 3T3-L1 cells impaired adipogenesis by direct cleavage of C/EBPβ mRNA and selected pre-miRNAs [9, 10].

In the present study, we demonstrated a lack of metabolic manifestation in Mcpip1^fl/fl^Alb^Cre^ mice, despite the deletion of Mcpip1 in hepatocytes. Although Mcpip1 knockout in hepatocytes reduces the expression of genes involved in glucose metabolism (*Pck1*, *G6pc*, *Irs1*), its concentration in the plasma was not changed when compared to Mcpip1^fl/fl^ counterparts. Similarly, the lack of Mcpip1 in hepatocytes did not influence lipid metabolism in mice fed chow or HFD. Notably, our observations are in accordance with another study, where we demonstrated that the deletion of Mcpip1 in Mcpip1^fl/fl^Alb^Cre^ mice is asymptomatic, in terms of blood morphology and inflammation markers detected in plasma, but leads to the massive hyperplasia of cholangiocytes and development of primary biliary cholangitis [Kotlinowski et al., submitted]. Recently, total Mcpip1 knockout mice were characterized by diminished hepatic triglyceride content [30], but as demonstrated here, this phenotype cannot be attributed solely to Mcpip1 depletion in hepatocytes, since dyslipidemia was detected only in Mcpip1^fl/fl^LysM^Cre^, but not in Mcpip1^fl/fl^Alb^Cre^ mice. What is more, the depletion of Mcpip1 in myeloid leukocytes led not only to dyslipidemia, but also to systemic proinflammatory phenotype resembling cachexia symptoms and bile ducts’ failure.

In our opinion, changes in lipid and glucose metabolism in Mcpip1^fl/fl^LysM^Cre^ mice are secondary to the inflammatory syndrome, rather than a direct product of myeloid leukocytes Mcpip1 activity on the control of hepatic metabolism. In fact, hypoglycemia and dyslipidemia are common features of murine cachexia models. Inflammatory syndrome and cachexia were already reported in Mcpip1^fl/fl^LysM^Cre^ animals by Li and coworkers [23]. Authors identified C/EBPβ and C/EBPδ, as targets of Mcpip1 RNase, being two critical transcriptional factors that drive cytokine production and lung injury. Moreover, mice with a myeloid-specific deletion of Mcpip1 were extremely sensitive to LPS-induced lung injury due to overproduction of proinflammatory cytokines and chemokines [23].

Cachexia is defined as a wasting syndrome, characterized by weight loss, systemic metabolic dysfunction, and immune system impairment [31, 32], all seen in Mcpip1^fl/fl^LysM^Cre^ mice. In accordance with literature data, Mcpip1^fl/fl^LysM^Cre^ animals were characterized by weight loss (13% from initial body weight). Although cachectic patients have normal serum levels of glucose [33], Mcpip1^fl/fl^LysM^Cre^ mice were hypoglycemic both after fasting or intraperitoneal glucose injection. They were even hypoglycemic upon feeding with a high fat diet, which in Mcpip1^fl/fl^ and Mcpip1^fl/fl^Alb^Cre^ mice induced glucose intolerance. Similarly to our mice, hypoglycemia was also observed in cancer cachexia murine model, namely Colon 26 (C26) adenocarcinoma xenograft model [34]. Hypoglycemia of Mcpip1^fl/fl^LysM^Cre^ mice might be explained by the altered expression of genes encoding proteins involved in gluconeogenesis (*G6pc*), insulin response (*Irs1, Irs2*), and glucose bidirectional transport (*Slc2a2*). Additionally, a raised level of Tnfα in plasma of these mice could account for decreased gluconeogenesis rates [35]. Also, dyslipidemia detected in Mcpip1^fl/fl^LysM^Cre^ mice is in line with results obtained from humans and C26 cachexia mouse model [34]. C26 mice are characterized by alterations in cholesterol metabolism reflected by low HDL and high LDL plasma level [36], seen also in Mcpip1^fl/fl^LysM^Cre^ mice. Studies of cachectic patients show, that their dyslipidemia results from a reduced HDL plasma level, which comes from anorexia [37], with a concomitant increase in LDL plasma level [38]. According to Pin et al., an increased level of LDL is dependent on HDL level reduction, due to the fact that LDL maturation requires the acquirement of cholesterol esters from HDL particles. If this process is disturbed, low-density, cholesterol-poor LDL particles lose their affinity to LDL receptors, and remain in circulation [36].

In Mcpip1^fl/fl^LysM^Cre^ mice we observed a very significant systemic inflammation, with some hepatic leukocyte infiltration, bile ducts pathology and extramedullary hematopoiesis (EMH) detected in the liver. The pathologic EMH, may be caused by one of many hematological diseases, bone marrow irradiation or anemia. [39, 40]. It can be also seen as part of the response to systemic inflammation or infection [39, 40]. In fact, both total and myeloid-specific Mcpip1 knockouts spontaneously develop systemic inflammatory response, leading to splenomegaly, lymphadenopathy and hyperimmunoglobulinemia [3, 13, 23]. Additionally, Kidoya and coworkers has shown recently, that post-transcriptional regulation mediated by Mcpip1 is essential for hematopoietic stem and progenitor cell homeostasis. Mcpip1 regulates self-renewal of hematopoietic stem and progenitor cells through modulating the stability of Gata2 and Tal1 mRNA and its dysfunction leads to the rapid onset of abnormal hematopoiesis [41].

Finally, Mcpip1^fl/fl^LysM^Cre^ mice did not develop hepatic steatosis, despite being fed a HFD. This finding is surprising since, it was reported earlier that cachectic patients and mice spontaneously develop hepatic steatosis [42]. Steatosis present in cachectic patients and animal models was caused, among others, by an enhanced free fatty acids uptake by hepatocytes, reduced β-oxidation, or by a high concentration of proinflammatory cytokines (e. g. Tnfa) in the plasma – all features present in Mcpip1^fl/fl^LysM^Cre^ mice [43–46]. However, as stated by Kliewer et.al, hepatic lipid metabolism may depend on the stage of cachexia, with no steatosis in early cachexia [47]. Cachexia in mice might be divided into an early or late stage, based on the difference in body weight between cachectic and control mice. As proposed by Sun and coworkers, this value was 18% for early and 29% for late cachexia [48]. In the case of Mcpip1^fl/fl^LysM^Cre^ mice, we observed a 21% difference in body weight between Mcpip1^fl/fl^ and Mcpip1^fl/fl^LysM^Cre^ that might suggest an early stage of cachexia, further supported by the lack of liver steatosis. We postulate, based on plasma biochemistry and liver qPCR data, that an uptake of circulating fatty acids in Mcpip1^fl/fl^LysM^Cre^ mice fed with high-fat diet might be reduced, explaining the lack of hepatic steatosis in these animals. Moreover, the expression of *Srebf* coding for the master regulator of lipogenesis in liver – transcription factor Sterol regulatory element-binding protein 1c – and its downstream targets [49] was reduced in the livers of Mcpip1^fl/fl^LysM^Cre^ mice.

In conclusion, we have demonstrated that depletion of Mcpip1 in myeloid leukocytes results in systemic inflammation, dyslipidemia, hypoglycemia and disturbed liver lipid metabolism all related with cachexia than direct effect of Mcpip1 knockout. On the other hand, Mcpip1^fl/fl^Alb^Cre^ mice did not exhibit signs of systemic inflammation or disturbed glucose and lipid metabolism.

## Abbrevations

ALT: alanine aminotransferase
AST: aspartate aminotransferase
BMI: body mass index
ECM: extracellular matrix
GTT: glucose tolerance test
HFD: high-fat diet
HDL: high-density lipoprotein
LDL: low-density lipoprotein
LDH: lactate dehydrogenase
NAFLD: non-alcoholic fatty liver disease
NASH: non-alcoholic steatohepatitis

## Data availability statement

Data can be shared upon request. For this purpose please contact Corresponding Author.

## Acknowledgements/grant support

This work was supported by research grant from National Science Centre, Poland no. 2015/19/D/NZ5/00254 to JeKo.

---

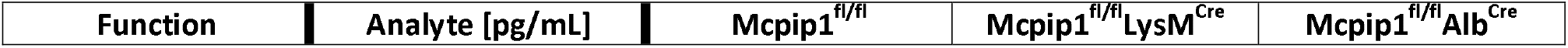

