## Supplemental data for "Role of Mcpip1 in obesity-induced hepatic steatosis as determined by myeloid and liver-specific conditional knockouts"

Jerzy Kotlinowski

Address: Gronostajowa Street 7, 30-387 Krakow, Poland

Table S1. Sequences of primers used for genotyping.

| **Locus** | **Forward** | **Reverse** |
| --- | --- | --- |
| **Loxp** | loxP: GCCTCTTGTCACTCCCTCCTCC | loxP-WT: GCCTTCCTGATCCTATTGGAG |
|  |  | loxP-mut: GAGATGGCGCAGCGCAATTAAT |
| **LysM-Cre** | LysM-Cre-WT: TTACAGTCGGCCAGGCTGAC | LysM-Cre: CTTGGGCTGCCAGAATTTCTC |
|  | LysM-Cre-mut: CCCAGAAATGCCAGATTACG |  |
| **Alb-Cre** | Alb-Cre-WT: TGCAAACATCACATGCACAC | Alb-Cre: TTGGCCCCTTACCATAACTG |
|  | Alb-Cre-mut: GAAGCAGAAGCTTAGGAAGATGG |  |

Table S2. Sequences of primers used for real-time PCR.

| **Gene** | **Species** | **Sequence** |
| --- | --- | --- |
| *Ef2* | Mus musculus | F: GACATCACCAAGGGTGTGCAG  R: TCAGCACACTGGCATAGAGGC |
| *Pck1* | Mus musculus | F: CGAGACTAGCGATGGGGGTG  R: CACATGGTTCCGCGTCCTG |
| *G6pc* | Mus musculus | F: CCGGATCTACCTTGCTGCTC  R: GCATTGTAGATGCCCCGGAT |
| *Irs1* | Mus musculus | F: CGGGTTCGAGAGAGCAGCAACA  R: CGGGTTCGAGAGAGCAGCAACA |
| *Irs2* | Mus musculus | F: GCCACAGTCGTGAAAGAGTGA  R: GTTGGTCGGAAACATGCCAA |
| *Slc2a2* | Mus musculus | F: GCTGCTGGATAAATTCGCCT  R: CGGGGTCCTTGGCTGAAAAA |
| *Cd36* | Mus musculus | F: GCAGCCTCCTTTCCACCTT  R: GGCATTGGCTGGAAGAACAA |
| *Fabp1* | Mus musculus | F: ACCCAAAGTGGTCCGCAATG  R: GACGACTGCCTTGACTTTTTCC |
| *Mttp* | Mus musculus | F: GCAGTTCTCACAGTACCCGT  R: TCCCCTGCCTGTAGATAGCC |
| *Ppara* | Mus musculus | F: TAATTTGCTGTGGAGATCGGC  R: AGCTTTGGGAAGAGGAAGGTG |
| *Acox1* | Mus musculus | F: ATGCCTTTGTTGTCCCTATC  R: CCATCTTCAGGTAGCCATTATC |
| *Cpt1a* | Mus musculus | F: AGGAGAATGCCAGGAGGTCA  R: GTGAGGCAGAACTTGCCCAT |
| *Acadl* | Mus musculus | F: ACGTCTGGACTCCGGTTCTG  R: TGCAATCGGGTACTCCCACA |
| *Srebf1* | Mus musculus | F: GGAACTTTTCCTTAACGTGGGC  R: ATGAGCTGGAGCATGTCTTCG |
| *Cebpb* | Mus musculus | F: CAAGCTGAGCGACGAGTACA  R: CAGCTGCTCCACCTTCTTCT |
| *Pparg* | Mus musculus | F: AGGCCGAGAAGGAGAAGCTGTTG  R: TGGCCACCTCTTTGCTCTGCTC |
| *Acc1* | Mus musculus | F: AATGAACGTGCAATCCGATTTG  R: ACTCCACATTTGCGTAATTGTTG |
| *Fasn* | Mus musculus | F: CTGCGGAAACTTCAGGAAATG  R: GGTTCGGAATGCTATCCAGG |
| *Scd1* | Mus musculus | F: TATCCTGGTTTCCCTGGGTGC  R: GGAACTCAGAAGCCCAAAGCTC |
| *Dgat2* | Mus musculus | F: GCCGATGGGTCCAGAAGAAG  R: CGATGTCTTTCTGGGTCGGG |
| *Il6* | Mus musculus | F: ACTTCACAAGTCGGAGGCTT  R: GGTACTCCAGAAGACCAGAGG |
| *Il1b* | Mus musculus | F: ACCCCAAAAGATGAAGGGCT  R: ACAGCTTCTCCACAGCCACA |
| *Tnfa* | Mus musculus | F: AGGCACTCCCCCAAAAGATG  R: GCTCCTCCACTTGGTGGTTT |
| *Cd68* | Mus musculus | F: GCTCCTCCACTTGGTGGTTT  R: AAGCCCCACTTTAGCTTTACC |
| *Tgfb* | Mus musculus | F: AGACCTTGAACCGCATCCTG  R: AGATGACAGCCTTCCCGTTG |

Table S3: Immune profile of plasma samples from 22-week-old mice fed with HFD for 12 weeks, determined by Luminex Assay. Out of 45 analytes, 14 has not been detected, namely: IL-2, IL-1β, IL-3, IL-7, CCL4/MIP-1 beta, CCL2/MCP1, RAGE, IL17e/IL25, IL17/IL17a, IL1a/IL-1F1, IL-4, IL-33, CCL5/RANTES, FGF-basic. For Mcpip1^fl/fl^ n=7, for Mcpip1^fl/fl^LysM^Cre^ n=7, for Mcpip1^fl/fl^Alb^Cre^ n=7. The table shows means ± SEM; * p<0.05; ** p<0.01; *** p<0.001 for Mcpip1^fl/fl^LysM^Cre^ or Mcpip1^fl/fl^Alb^Cre^ vs Mcpip1^fl/fl^; # p<0.05 for Mcpip1^fl/fl^LysM^Cre^ fed HFD vs Mcpip1^fl/fl^LysM^Cre^ fed control diet, nd – not detected.

| **Function** | **Analyte [pg/mL]** | **Mcpip1^fl/fl^** | **Mcpip1^fl/fl^LysM^Cre^** | **Mcpip1^fl/fl^Alb^Cre^** |
| --- | --- | --- | --- | --- |
| **Interleukins** | **IL-5** | nd | 0.043 ± 0.006 *** | nd |
|  | **IL-6** | nd | 0.020 ± 0.006 ** | nd |
|  | **IL-6 R alpha** | 11.22± 0.758 | 26.55 ± 1.299 *** | 12.01 ± 0.586 |
|  | **IL-10** | nd | 0.024 ± 0.004 *** | nd |
|  | **IL-12 p70** | nd | 0.084 ± 0.075 | nd |
|  | **IL-13** | nd | 0.191 ± 0.030 *** | nd |
|  | **IL-16** | 0.473 ± 0.082 | 0.781 ± 0.145 | 0.446 ± 0.046 |
|  | **IL-27** | nd | 0.031 ± 0.012 * | nd |
| **Chemokines** | **CCL8/MCP-2** | 104.28 ± 8.57 | 605.85 ± 56.12*** | 102.11 ± 4.70 |
|  | **CCL7/MARC** | 0.048 ± 0.004 | 0.191 ± 0.037 ** | 0.061 ±0.013 |
|  | **CCL12/MCP-5** | 0.081 ± 0.049 | 0.136 ± 0.035 | 0.034 ± 0.004 |
|  | **CCL3/MIP-1 alpha** | 1.02 ± 0.11 | 11.51 ± 3.50 ** p=0.061 vs AIN | 0.64 ±0.09 |
|  | **CCL20/MIP-3 alpha** | 0.055 ± 0.036 | 0.082 ± 0.039 | 0.027 ± 0.027 |
|  | **CCL19/MIP-3 beta** | 0.0004 ± 0.0002 | 0.028 ± 0.007 *** | nd |
|  | **CCL11/Eotaxin** | 1.44 ± 0.10 | 1.22 ± 0.21 | 1.18 ± 0.06 |
|  | **CXCL16** | 0.274 ± 0.024 | 0.199 ± 0.058 | 0.246 ± 0.018 |
| **Growth factors** | **GM-CSF** | 0.0006 ± 0.0002 | 0.0005 ± 0.0005 | 0.00009 ± 0.00006 |
|  | **M-CSF** | 3.61 ± 0.40 | 2.30 ± 0.86 | 5.22 ± 0.80 |
|  | **G-CSF** | 0.102 ± 0.020 | 0.159 ± 0.032 | 0.139 ± 0.033 |
| **TNF** | **TNF-alpha** | 2.95 ± 0.01 | 3.80 ± 0.26 ** | 2.95 ± 0.02 |
|  | **TNF RI** | 0.691 ± 0.019 | 1.041 ± 0.038 *** | 0.659 ± 0.042 |
|  | **TNF RII** | 2.36 ± 0.42 | 7.66 ± 0.91 *** | 3.47 ± 0.08 |
| **Angio-genesis** | **VEGF** | nd | 0.004 ± 0.001 ** | nd |
|  | **VEGF R2** | 40.04 ± 1.65 | 23.94 ± 1.53 *** | 46.69 ± 1.87 |
|  | **Endoglin** | 2.51 ± 0.36 | 1.35 ± 0.14 * | 2.28 ± 0.18 |
| **Others** | **Trombospondin-4** | 664.99 ± 78.20 | 278.84 ± 35.49 ** | 617.93 ± 24.54 |
|  | **CHI3-L1** | 34.45 ± 1.60 | 268.74 ± 38.21 *** # | 32.18 ±2.33 |
|  | **Icam-1** | 11.32 ± 0.86 | 16.07 ± 2.68 | 10.79 ± 1.73 |
|  | **p-Selectin/CD-62P** | 27.88 ± 1.69 | 53.52 ± 5.62 *** | 30.45 ± 0.67 |
|  | **Syndecan-1/CD138** | 5.63 ± 0.36 | 40.91 ± 6.78 *** # | 5.84 ± 0.34 |
|  | **IFN gamma** | 0.0320 ± 0.0001 | 0.0330 ± 0.0006 | 0.03200 ± 0.00006 |

**Figure S1: Comparison of Mcpip1^fl/fl^ controls from Mcpip1^fl/fl^LysM^Cre^ and Mcpip1^fl/fl^Alb^Cre^ strains.**

All data obtained from Mcpip^fl/fl^ mice were separated depending of the strain of origin. For mice fed with control diet: (A) Body/liver ratio, level of cholesterol, LDL, HDL in plasma and AUC; (B) Expression of selected genes in liver: *Slc2a2*, *Fabp1*, *Ppara*, *Srebf1*, *Tnfa*, *Mmp3*; (C) Level of analytes in plasma samples determined by Luminex Assay: IL-6 R alpha, CCL8/MCP-2, G-CSF, Tnf-alpha, Vegf-R2, Syndecan-1/CD138.

For mice fed with HFD: (D) Body/liver ratio, level of cholesterol, LDL, HDL in plasma and AUC; (E) Expression of selected genes in liver: *Slc2a2*, *Fabp1*, *Ppara*, *Srebf1*, *Tnfa*, *Mmp3*; (F) Level of analytes in plasma samples determined by Luminex assay: IL-6 R alpha, CCL8/MCP-2, G-CSF, Tnf-alpha, Vegf-R2, Syndecan-1/CD138.

For Mcpip^fl/fl^ all n=10 (control diet: Body/liver ratio, Cholesterol, LDL, HDL), n=8 (control diet: AUC, gene expression), n=6 (control diet: Luminex); n=7 (HFD: Body/liver ratio, Cholesterol, LDL, HDL, AUC, gene expression, luminex); for Mcpip1^fl/fl^ control for Mcpip1^fl/fl^LysM^Cre^ n=5 (control diet: Body/liver ratio, Cholesterol, LDL, HDL), n=4 (control diet: AUC, gene expression; HFD: luminex), n=2 (control diet: Luminex; HFD: Body/liver ratio, Cholesterol, LDL, HDL); n=3 (HFD: AUC, gene expression); for Mcpip1^fl/fl^ control for Mcpip1^fl/fl^Alb^Cre^ n=5 (control diet: Body/liver ratio, Cholesterol, LDL, HDL; HFD: Body/liver ratio, Cholesterol, LDL, HDL), n=4 (control diet: AUC, gene expression, Luminex; HFD: AUC, gene expression), n=3 (HFD: luminex). The graphs show means ± SEM.

**Figure S2: Daily food intake, ALT and AST activity and liver mass of Mcpip1^fl/fl^, Mcpip1^fl/fl^LysM^Cre^ and Mcpip1^fl/fl^Alb^Cre^ mice.**

Daily food intake was analyzed at the age of 12, 16 and 20 weeks for chow food (A) and HFD (F). (B,G) Liver mass and (C,H) liver/body ratio. Plasma analysis of 22-week-old mice: (D,I) ALT, (E,J) AST. For Mcpip1^fl/fl^ n=7 (A,F,I,J), n=10 (B,C,D,E,G,H); for Mcpip1^fl/fl^LysM^Cre^ n=6 (A,D,E,F,I,J), n=4 (B,C), n=10 (G,H); for Mcpip1^fl/fl^Alb^Cre^ n=6 (A,F,I,J), n=8 (D,E), n=10 (B,C,G,H). The graphs show means ± SEM; * p<0.05; ** p<0.005; *** p<0.001.
