## Supplementary figures and images for "Role of Mcpip1 in obesity-induced hepatic steatosis as determined by myeloid and liver-specific conditional knockouts"

### Supplemental Figure 1

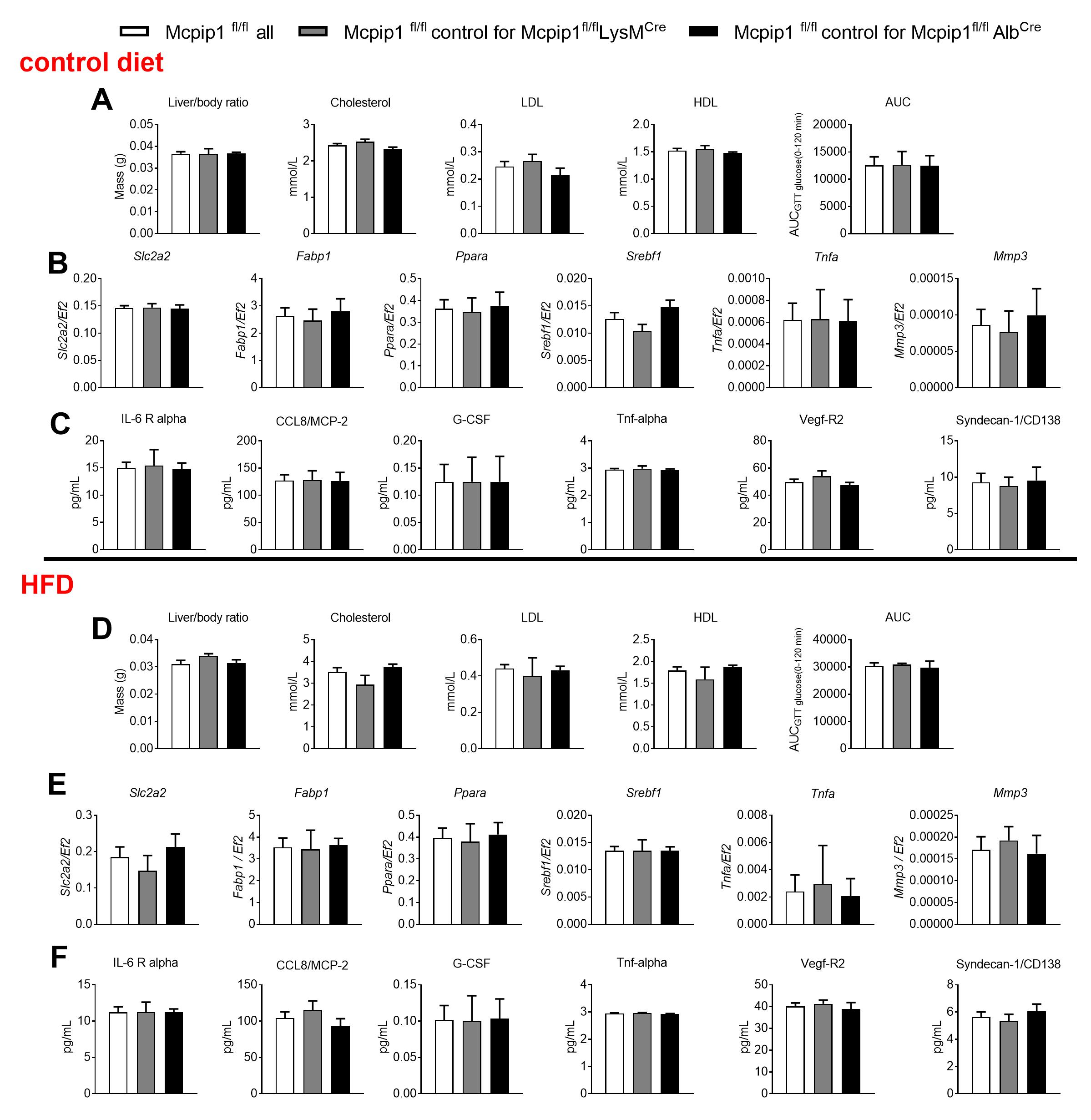

### Supplemental Figure 2

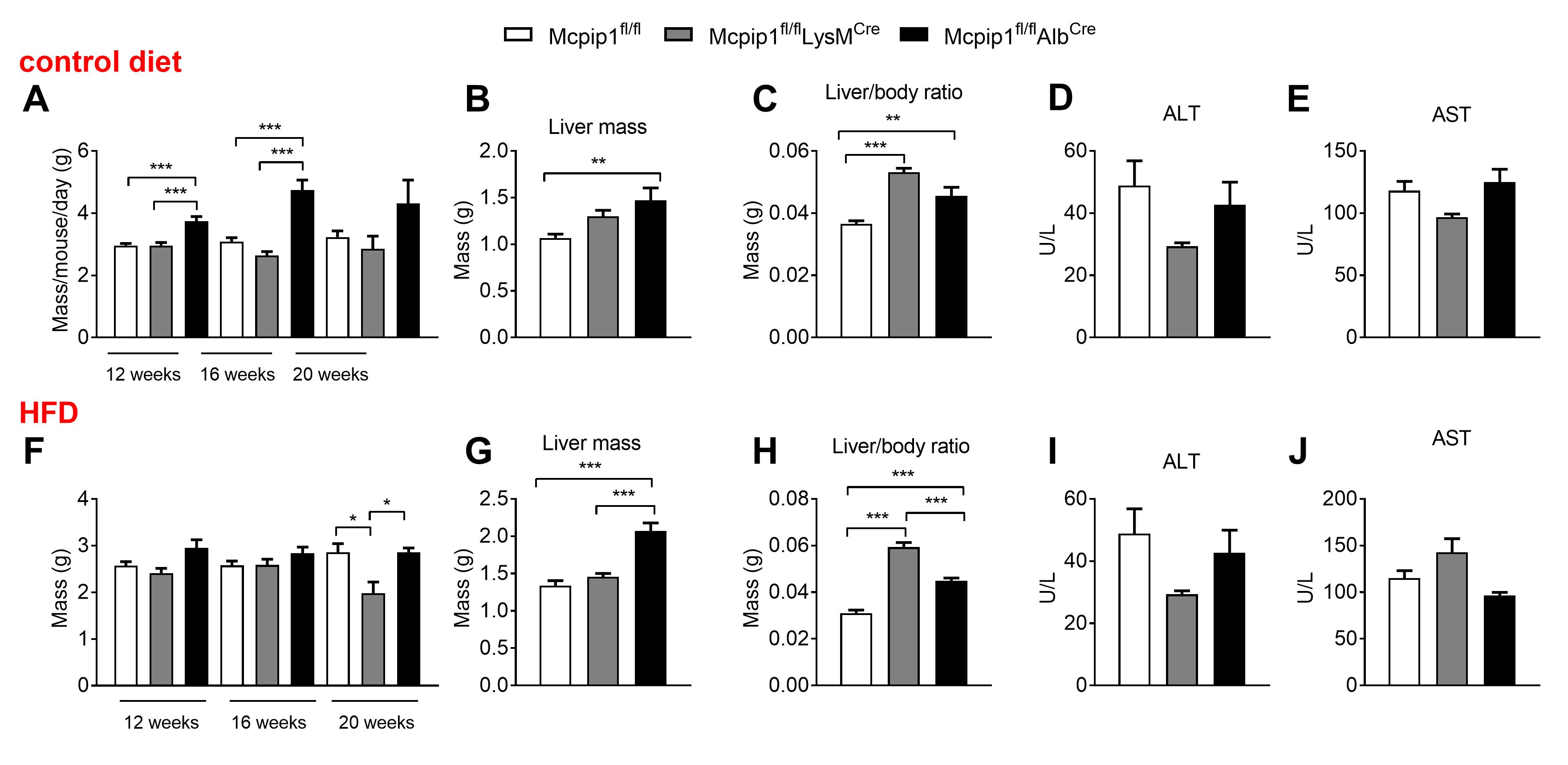
